# Antivirals for Monkeypox Virus: Proposing an Effective Machine/Deep Learning Framework

**DOI:** 10.1101/2024.02.11.579829

**Authors:** S. Morteza Hashemi, Arash Zabihian, Masih Hajsaeedi, Mohsen Hooshmand

## Abstract

Monkeypox is one of the infectious viruses which caused morbidity and mortality problems in these years. Despite its danger to public health, there is no approved drug to stand and handle Monkeypox. On the other hand, drug repurposing is a promising screening method for the low-cost introduction of approved drugs for emerging diseases and viruses which utilizes computational methods. Therefore, drug repurposing is a promising approach to suggesting approved drugs for the monkeypox virus. This paper proposes a computational framework for monkeypox antiviral prediction. To do this, we have geenrated a new virus-antiviral dataset. Moreover, we applied several machine learning and one deep learning method for virus-antiviral prediction. The suggested drugs by the learning methods have been investigated using docking studies. To the best of our knowledge, this work is the first work to study deep learning methods for the prediction of monkeypox antivirals. The screening results confirm that Tilorone, Valacyclovir, Ribavirin, Favipiravir, and Baloxavir marboxil are effective drugs for monkeypox treatment.

## 1 Introduction

Monkeypox (MPXV) is a viral zoonosis disease resulting from an enveloped double-stranded DNA virus that belongs to the Poxviridae family and causes an international public health emergency [1]. The initial occurrence of monkeypox in animals and humans was reported in 1958 and 1970, respectively [2]. A global epidemic of MPXV in May 2022 resulted in more than 85,000 cases across regions with no history of transmission [3]. Also, over 30,000 cases of MPXV had been reported in the US as of March 2023 [4]. The virus transmits through contact with skin lesions, respiratory droplets, body fluids, and fomites [2]. Although several antiviral drugs are suggested for treatment against MPXV, the Food and Drug Administration (FDA) has not approved any specific drugs for human monkeypox [5]. On the other hand, drug repurposing is an efficient approach for accelerating the discovery of novel treatments by providing new indications for approved drugs [6]. Currently, modern computational methods are making a significant impact on drug repurposing for the treatment of viral infectious diseases. For example, drug repurposing has played an important role in research and proposing novel drugs for Covid 19. Tang et al. suggested a non-negative matrix factorization method for suggesting antivirals against SARS-CoV-2 [7]. However, the matrix factorization methods suffer from two problems, lack of generalizability and data leakage [8]. Beck et al. SARS-CoV-2 [9] using deep model. Zeng et al. have the same mission to suggest new drugs for SARS-CoV-2 using deep models [10]. Other studies suggested and showed the use of machine learning and a spectrum of deep learning methods have a higher performance and more trustful results for drug-target prediction of proteins, SARS-CoV-2, and other viruses [9, 11–19]. It is worth mentioning that although deep learning algorithms are the winners in the field of drug repurposing, the machine learning methods can reach a similar performance in most cases with a lower amount of resource consumption [20]. Therefore, this paper tends to check and propose methods from both machine learning and deep learning for monkeypox antiviral prediction.

This study is centered on using computational drug repurposing to identify a novel candidate for targeting monkeypox among approved antiviral drugs. To do this, we have created a virus-antiviral dataset and gathered similar viruses to the MPXV. In addition to that, we apply proper *similarity measures* to create effective features for viruses and antivirals. We, as mentioned above, utilize machine learning and deep learning methods for the prediction.

Then, based on the results of the learning methods, we chose the most promising drugs and performed a docking study to approve the results of the machine learning methods. The docking study confirms that the results of the learning methods are promising and proper voices for MPXV treatment. To the best of our knowledge, this is the first work in applying results of deep and machine learning for drug recommendation for the MPXV virus.

The contribution of the paper is 5-fold.

- Generating a virus-antiviral dataset for MPXV prediction.
- Applying machine learning algorithms, i.e., decision tree, SVM, and random forest algorithms.
- Proposing and applying a CNN model to train the prediction model.
- Proposing five approved drugs for MPXV treatment, i.e., *Tilorone, Valacyclovir, Ribavirin, Favipiravir, and Baloxavir marboxil*.
- Validating the proposed drugs by docking study.

The results are promising and the docking study supports the proposed MPXV antivirals with high scores. The structure of the paper is as follows. Section 2 explains the dataset generation and the learning methods for the MPXV prediction. Additionally, it introduces the docking studies applied to the proposed antivirals. Section 3 compares the performance of learning algorithms, introduces the proposed drugs, and reports docking studies on the proposed drugs. Section 4 finalizes the paper.

## 2 Materials and methods

This paper proposes a prediction framework for monkeypox virus-antiviral interaction that uses computational learning methods and we call it *MPXV-Pred*. This section provides the steps of MPXV-Pred. The whole process contains five main steps, i.e., data collection, dataset generation, the learning phase using machine learning and deep learning methods, the voting procedure to propose the most effective antiviral, and finally the docking step. Figure 1 shows the pipeline of the proposed method. The following sections describe these steps.

**Fig 1.**
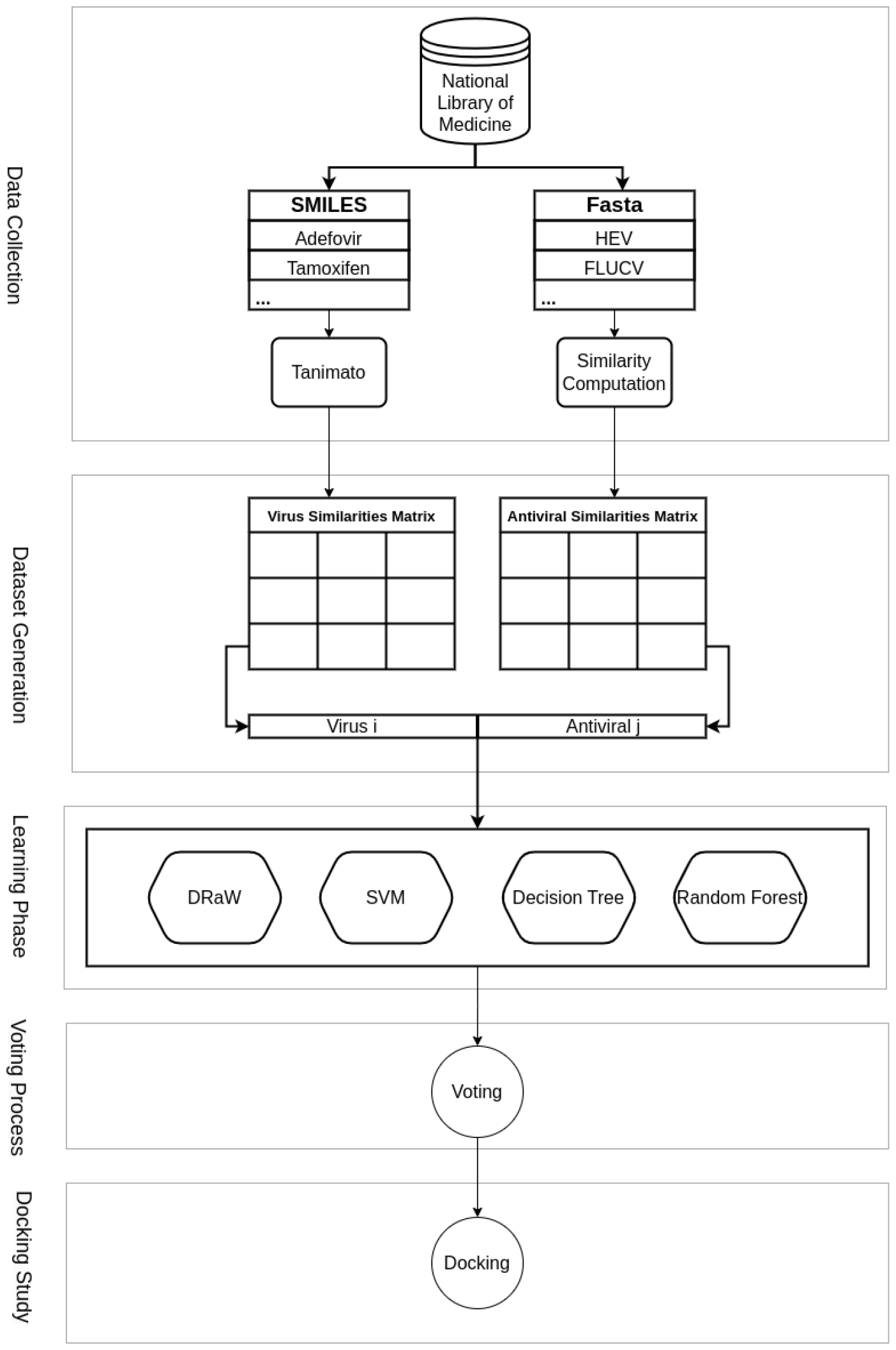
MPXV-Pred Framework. It aims to predict novel antivirals for the MPXV. The pipeline contains five steps data collection, dataset generation, machine and deep learning methods, the voting procedure to announce the promising antiviral, and docking study. I) Data collection: the raw and primary information on viruses and antivirals have been collected from the *NIH*. The approved virus-antiviral interactions have also been provided from DrugVirus.info 2 [21]. II) Dataset generation: the output of this step is the generated dataset including the similarity matrix of viruses, similarity matrices of antivirals, and virus-antiviral interaction matrix. The virus similarity matrix has been computed using *sequence alignment* and calculating the similarity percent. Additionally, the Tanimoto score has been used to compute the antiviral similarity matrix. III) The learning phase: this phase employs four different machine learning methods, i.e. CNN, SVM, Decision tree, and Random forest. IV) The voting step uses the prediction results of the ML methods of the previous step and votes among them to suggest the efficient antiviral for MPXV. V) The Docking study investigates and approves the predicted antivirals for MPXV treatment.

### 2.1 Data collection

There exists no proper dataset available to contain information on viruses and antivirals that have common properties with the MPXV virus. Therefore, in the first step, it is necessary to prepare a new corresponding dataset. To do that we collected raw data of viruses and antivirals, e.g., smallpox from “The National Library of Medicine” databases [22]. The raw data includes the Fasta format of virus sequences of 100 viruses, the simplified molecular-input line-entry system (SMILES) format of 198 drugs, and the interaction between them. The collected dataset has 96 approved virus-antiviral interactions, therefore, its sparsity reaches 99.52%.

### 2.2 Similarity computation and generating the dataset

Using sequences and SMILES, the similarities between each pair of viruses in addition to each pair of drugs were computed correspondingly. Tanimoto [23] is a popular approach to calculating the similarities between drugs. Thus, we applied this algorithm to the collected data to create a similarity matrix of drugs. To calculate the similarity among the viruses we align them in the first step. Therefore, we use the PairWiseAligner tool from the BioPython library [24]. The alignments are performed locally with the Smith-Waterman algorithm [25] and the NUC44 substitution matrix [] is used to score the alignments. The gap score is set to−10 and the gap extend score is set to −0.5. These two scores are default scores from emboss [26] tools. Figure 1-a shows the procedure of collecting the data and inverting it to similarity matrices.

### 2.3 Learning phase

The learning phase aims to train a model to predict the most effective antivirals for the MPXV. To do this, MPXV-Pred utilizes three machine learning methods, i.e., decision tree, SVM, and random forest [27]. The MPXV-Pred uses the similarity vectors of each virus and antiviral as their feature vector. Therefore, the input vectors of each learning method are the concatenation of feature vectors corresponding to each virus-antiviral pair. The interaction matrix plays the role of the virus-antiviral label. Therefore, we can formulate the problem as follows.

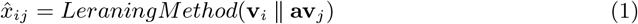

Where, **v**_*i*_ and **av**_*j*_ represent the features vectors of *i*-th virus and *j*-th antiviral. The 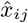 shows the prediction result of the interaction between the *i*-th virus and *j*-th antiviral.

Moreover, it uses a convolution-based deep learning model which we call our proposal as **D**rug **R**epurposing-**a**nalytic **W**ay (DRaW) [8] to complete the same mission. Figure 2 shows the DraW framework. DRaW consists of three convolution layers in total. The first layer is the convolution layer with 128 filters with size 3 and batch normalization is applied to improve training stability. And dropout layer is used to randomly drop 50% of the unit during training to reduce overfitting. for the second and third layers, we’ve used the same pattern but with different sizes of filters. And lastly, a dense layer with 128 units is applied for classification.

**Fig 2.**
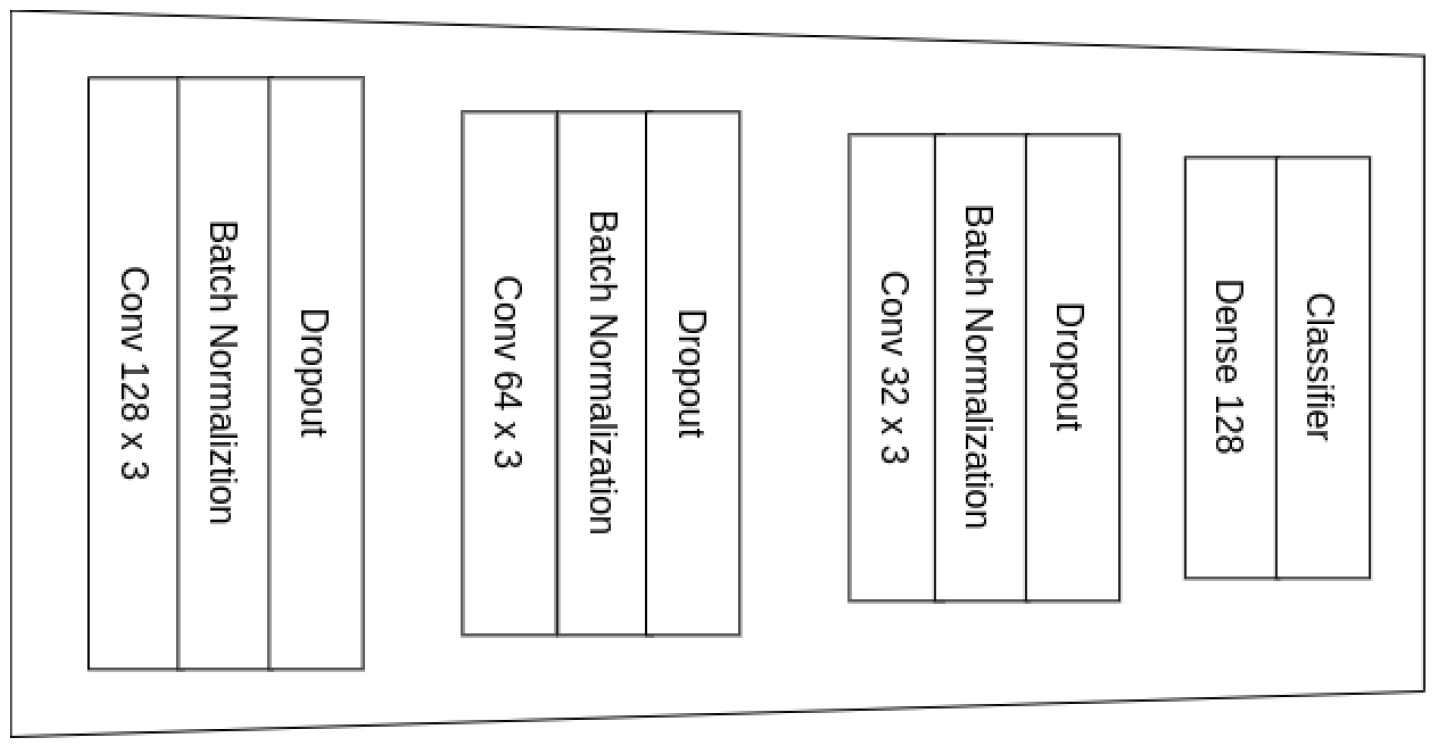
DRaW architecture. It is a CNN-based deep learning method.

DRaW’s input vector is the virus-antiviral pair’s representation from the concatenation of **v**_*i*_ and **av**_*j*_, or **e**_*i,j*_ = **v**_*i*_ ∥ **av**_*j*_.

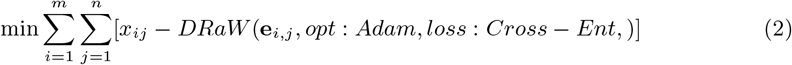

#### Voting process

The MPXV-Pred investigates the suggested antivirals of whole methods and reports those that happen in *at least two of the learning methods*. This may help to suggest the most effective potential antivirals for MPXV treatment.

### 2.4 Homology modeling and molecular docking studies

Structure-based molecular docking is a computational technique to evaluate how a ligand interacts with a target and addresses three main objectives: virtual screening, posture prediction, and estimation of binding affinity [28]. Since the outbreak of the MPXV global epidemic, *tecovirimat* is the only currently available therapeutic agent that has been suggested for use in Europe under “exceptional circumstances” by the European Medicines Agency [29]. In other words, there is no approved drug for MPXV. According to the literature, the envelope receptor F13 is a potential target for tecovirimat. This protein plays a crucial role in the growth and maturation of the monkeypox virus. The mechanism of F13-inhibition could be a promising treatment for monkeypox. Therefore, we employed molecular docking simulations to investigate the binding capability of drugs predicted by our proposed model to the F13 protein as a potential and promising target. Among the top ten drugs predicted by DRaW, we investigated instances that were also included in the predicted drug lists of three other models. To elucidate further, according to the data from Table 4, Tilorone, Valacyclovir, Ribavirin, Baloxavir, and Favipiravir are the drugs that appear most frequently across the predicted lists of all models. Therefore, we decided to do molecular docking studies employing these drugs. The primary challenge arises from the unavailability of the complete crystal structure of the poxvirus F13. To address this challenge, the UniprotKB [30] entry Q5IXY0, representing the monkeypox envelope protein F13 with a length of 372 amino acids, was uploaded to AlphaFold2 [31] to generate a predicted 3D protein structure. We also determined the predicted local distance difference test (pLDDT) values for the predicted structure. This modeled structure was optimized by energy minimization using the YASARA server [32], which used the YASARA force field for this purpose. The homology-modeled protein structure was prepared using the Autodock tools (ADT). This procedure involved the incorporation of polar hydrogens and Kollman charges. The 3D-SDF structures of all five candidate drugs were retrieved from NCBI PubChem [33]. The preparation of ligands involved the addition of polar hydrogens and Gasteiger charges. Additionally, root detection and selection of torsions from the torsion tree were conducted to rotate all the rotatable bonds [34]. We identified the F13 binding site based on the methodology outlined by Li et al. [35]. According to their study, the grid box was fixed at (−6.27702) ∗ (−2.48567) ∗ (−8.66086) in XYZ-coordinates, with a radius of 8.9, and the spacing for the grid box was kept at 0.375 Å. Finally, docking studies were performed by AutoDock 4.2 using the Lamarckian genetic algorithm.

### 2.5 Complexity analysis

We provide a complexity analysis of all four learning methods. Let *m* be the number of viruses and *n* be the number of antivirals. The feature vector is the concatenation of virus and antiviral similarity vectors. We assume its size is *d*.

#### Decision tree

Time complexity of the decision tree is *O*(*m ∗ n ∗ log*(*m ∗ n*) *∗ d*) where *m ∗ n* is the number of whole virus-antiviral pairs in the training phase and d is the dimensionality of the data. The runtime complexity is *O*(*depth*).

#### SVM

Train complexity of Support Vector Machine is *O*((*m ∗ n*)^2^). The runtime complexity is *O*(*k ∗ d*) where k is the number of support vectors and d is the dimensionality of data.

#### Random forest

The time complexity of a random forest depends on the number of trees in the forest and the depth of each tree. So the training time complexity is *O*(*m ∗ n ∗ log*(*m ∗ n*) *∗ d ∗ t*). Where t is the number of decision trees. The runtime time complexity is *O*(*depth ∗ t*).

#### Deep learning

We assume the number of epochs in the training phase is *e* and each epoch time interval is equal to *T*_*t*_. Then, the training phase time complexity is *O*(*m ∗ n ∗ d ∗ e ∗ T*_*t*_). Its runtime complexity is *O*(*d ∗ T*), where *T* is the test phase running time.

## 3 Results

This section provides the result of the MPXV-Pred using the proposed methods with early stpping. The reported results are based on 10-fold cross-validation.

Table 1 reports the specifications of the SVM, random forest, decision tree, and DRaW.

**Table 1.** Learning models’ parameters.

| ML Algorithms | Parameters | Parameters' Values |
| --- | --- | --- |
| DRaW | optimizer | adam |
|  | loss | BinaryCrossEntropy |
|  | epoch | 500 |
|  | batch size | 128 |
| SVM | kernel | rbf |
|  | degree | 3 |
|  | gamma | scale |
|  | tol | 1e-3 |
| Random Forest | criterion | gini |
|  | no. estimators | 100sqrt |
|  | max features | sqrt |
| Decision Tree | criterion | gini |
|  | splitter | best |

### 3.1 Performance evaluations

We evaluate the results of the methods using the following metrics. These metrics are useful for binary classification [36]. The MPXV-Pred dataset is imbalanced, therefore, metrics such as F1-score and AUPR are significant in interpreting the results [37].

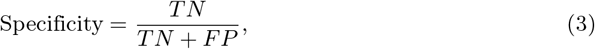

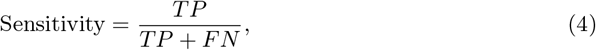

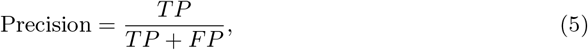

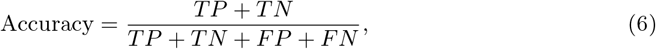

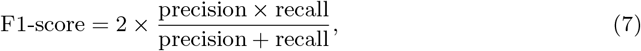

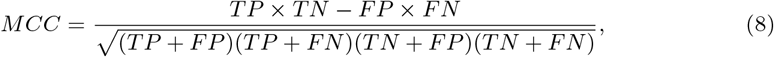

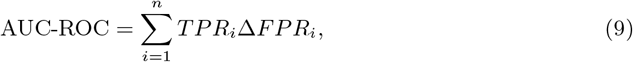

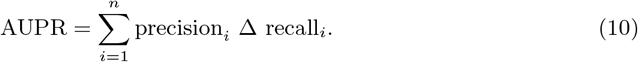

### 3.2 Prediction results

Table 2 and 3 show two different handling of the results. As mentioned above, we applied three machine learning methods and DRaW, a deep convolutional learning method for drug repurposing, to the dataset. Table 2 shows the result by defining a fixed threshold equal to 0.5. In the majority of metrics DRaW model has the highest score. DRaW has the highest AUC-ROC. Additionally, The highest AUPR score belongs to the decision tree and DRaW. We believe the higher result of the decision tree model in comparison to the random forest and SVM could be due to overfitting [38]. Because decision trees have a high probability of falling into the overfitting problem.

**Table 2.** Comparing the performance of machine learning models by different measures. In this experiment, the threshold limit in the classification problem is set equal to 0.5.

| Method | Sp | Re(Se) | Pr | Ac | F1 | MCC | AUC-ROC | AUPR |
| --- | --- | --- | --- | --- | --- | --- | --- | --- |
| Decision Tree | 0.9982 ± 0.0006 | 0.3337 ± 0.0324 | 0.4953 ± 0.0931 | 0.9949 ± 0.0006 | 0.3962 ± 0.0424 | 0.4028 ± 0.0479 | 0.6660 ± 0.0162 | 0.4161 ± 0.0536 |
| SVM | 0.9996 ± 0.0004 | 0.0816 ± 0.0387 | 0.5600 ± 0.2881 | 0.9951 ± 0.0003 | 0.1383 ± 0.0620 | 0.2062 ± 0.0875 | 0.8593 ± 0.0730 | 0.2456 ± 0.1060 |
| Random Forest | 1.0 ± 0.0 | 0.1329 ± 0.0283 | 1.0 ± 0.0 | 0.9957 ± 0.0004 | 0.2337 ± 0.0436 | 0.3622 ± 0.0379 | 0.9543 ± 0.0196 | 0.3890 ± 0.0408 |
| DRaW | 0.9999 ± 0.0002 | 0.1143 ± 0.0557 | 0.9200 ± 0.1789 | 0.9955 ± 0.0011 | 0.1966 ± 0.0842 | 0.3104 ± 0.0716 | 0.9641 ± 0.0172 | 0.4114 ± 0.0534 |

**Table 3.**
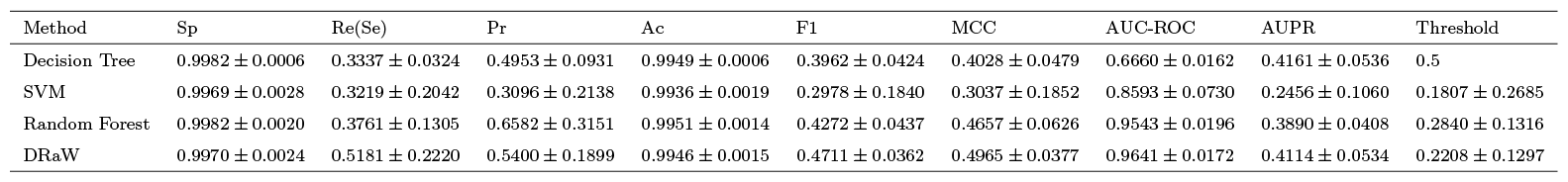
Comparing the performance of machine learning models by different measures. Since the dataset is imbalanced and only about 0.4% of the samples interact, it is regular to define the threshold limit as floating, unlike the previous test. The result of this experiment is shown in this table.

**Table 4.** Proposed drugs from the voting process.

| Draw | SVM | Random Forest | DecisionTree |
| --- | --- | --- | --- |
| Tilorone | Tilorone | Tilorone | Tilorone |
| Valacyclovir | Amantadine | Favipiravir | Ribavirin |
| Ribavirin | Baloxavir marboxil | Ribavirin |  |
| Foscarnet | Lamivudine | Itraconazole |  |
| Inosine (Inosine pranobex) | Favipiravir | Baloxavir marboxil |  |
| Baloxavir marboxil | Valacyclovir | Valacyclovir |  |
| Favipiravir | Thymalfasin | Fluvastatin |  |
| Valganciclovir | Rimantadine | Famciclovir |  |
| Lamivudine | Lopinavir | Ilaprazole |  |
| Ganciclovir | Zanamivir | Resveratrol |  |

In contrast to the previous calculation of evaluation metrics, we defined a floating threshold to choose the best values of all metrics except AUC-ROC and AUPR (Because these two do not depend on a single threshold). Table 3 shows the results of the floating threshold. The last row shows the threshold of each method. The decision tree model does not need any threshold and, therefore, is left empty. The DRaW has the highest AUC-ROC and the decision tree has the lowest AUC-ROC. The DRaW and decision tree have the highest AUPR. The high value of AUPR for the decision tree, as mentioned before, could be due to overfitting.

Figure 3 shows the bar chart of the AUC-ROC and AUPR of the decision tree, SVM, random forest, and DRaW models. As shown in the figure, random forest and DRaW have the highest AUC-ROC, and the winner is the latter. Additionally, DRaW and decision tree have the highest AUPR.

**Fig 3.**
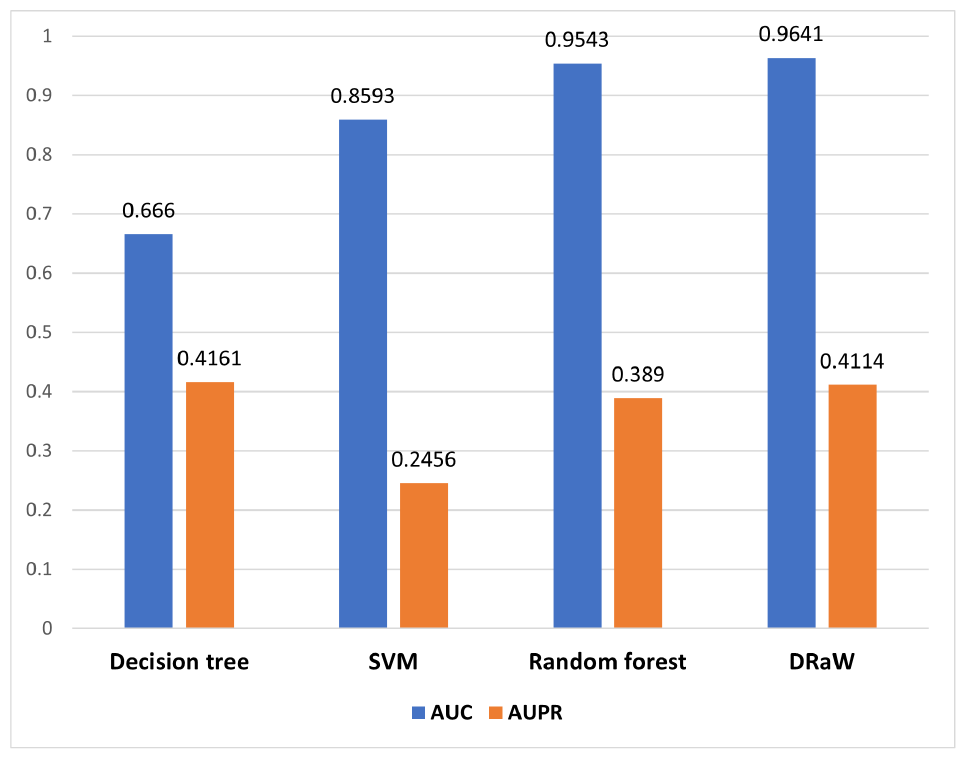
Performance comparison of methods on MPXV-Pred dataset. AUC-ROC and AUPR bar charts.

### 3.3 Voting results

Table 4 shows the suggested drugs of voting results of whole four methods. As shown in the table, the first rank in all methods belongs to Tilirone. Valacyclovir occupies the second rank in CNN and has been suggested by three out of four methods. Ribavirin has been suggested by three out of four methods. The same as the latter happened for Favipiravir. Finally, Baloxavir marboxil has been reported by three out of four methods. Notably, the decision tree suggested two drugs and we report those two, i.e., Tilirone and Ribavirin. One important observation is that while the four methods follow different approaches for learning and prediction, they confirm each other and the majority of top-rank antivirals are the same in all of them.

### 3.4 Homology modeling and molecular docking Results

We utilized AlphaFold tools to predict the three-dimensional structure of the monkeypox envelope protein F13, which has a length of 372 amino acids. The evaluation of structure prediction relied on the predicted local distance difference test (pLDDT) score [30]. The highest pLDDT score among the five predicted envelope protein F13 structures was 91.04. The additional pLDDT scores are shown in Table 5.

**Table 5.** pLDDT scores.

| Model Rank | Mean pLDDT | Max PAE | pTM |
| --- | --- | --- | --- |
| 1 | 91.04 | 30.95 | 0.92 |
| 2 | 90.67 | 30.77 | 0.92 |
| 3 | 90.26 | 30.86 | 0.92 |
| 4 | 90.21 | 31.13 | 0.91 |
| 5 | 89.41 | 30.81 | 0.91 |

Figure 4 Illustrates the homology model confidence score with predicted LDDT and predicted aligned error (Rank 1). Given that a pLDDT score greater than 70 is indicative of a high-quality structure, the results from the pLDDT analysis further confirm the high accuracy of the predicted structures.

**Fig 4.**
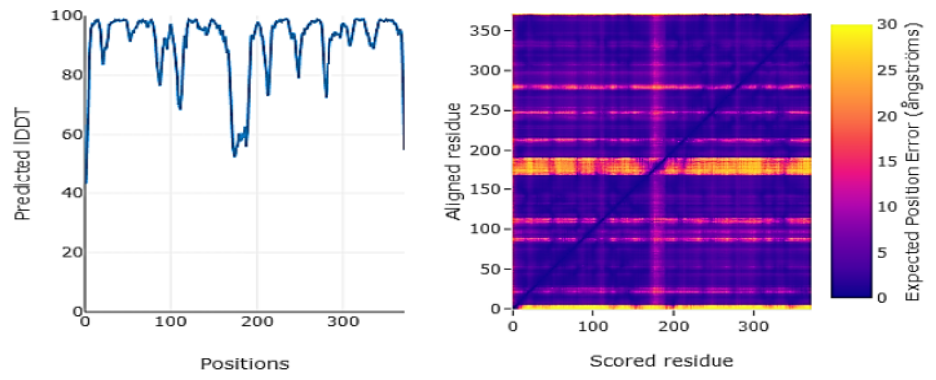
The confidence score of the homology model, along with the predicted Local Distance Difference Test (LDDT) and the predicted aligned error.

Molecular docking studies were conducted to assess the potential interactions between the F13 protein and a set of five chosen antiviral drugs. We chose the best ligand conformations by clustering them based on both RMSD and binding affinity [39]. To explore the intermolecular interaction forces, the results of the docking were visualized using Biovia Discovery Studio Visualizer [40]. Table 6 displays the five selected antiviral drugs along with their respective docking scores with the homology-modeled F13 protein structure. Based on the docking scores, Baloxavir exhibited the most favorable binding energy of -8.28 kcal/mol and formed three hydrogen bonds with LYS 88, ASN 90, and SER 58. Figures 1 to 5 illustrate both the 3D and 2D representations of each drug-F13 protein complex.

**Table 6.**
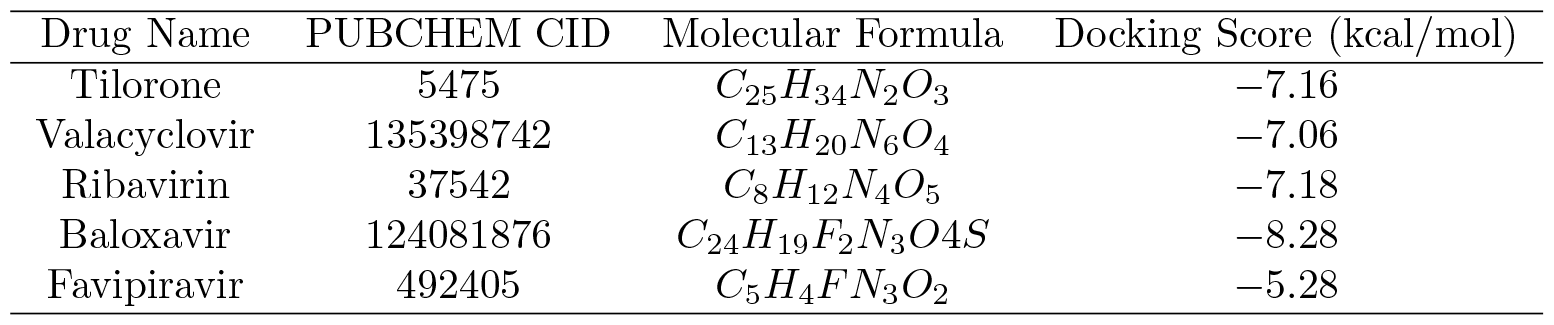
Docking scores of selected drugs.

Figure 5 shows Tilorone’s 3D and 2D representations. In this figure, the docking results reveal hydrogen bonds with ASP109 and ILE108 residues, as well as van der Waals interactions with several other residues. Additionally, the binding of tilorone to the F13 protein indicates unfavorable acceptor-acceptor interactions with HIS338.

**Fig 5.**
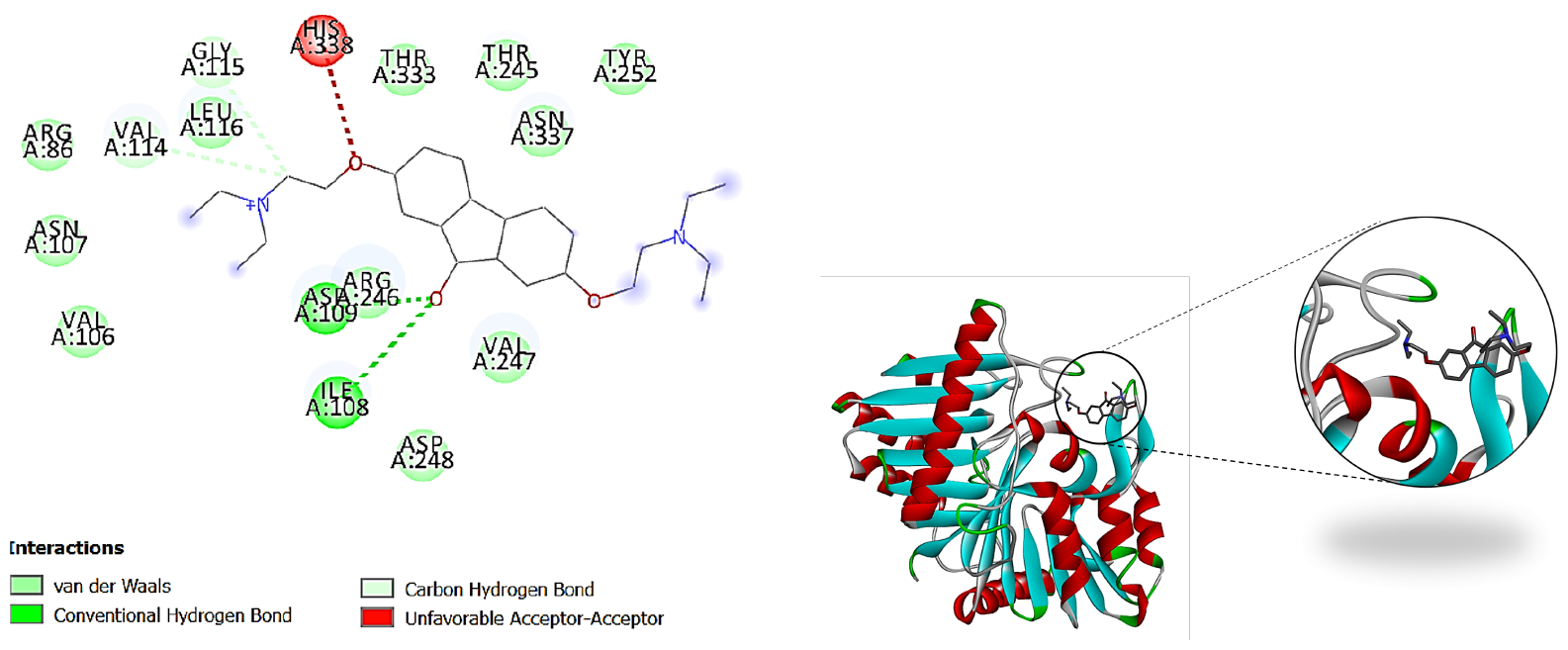
2D and 3D representations of the docked pose for the predicted interaction between Tilorone. The green dashed lines represent hydrogen bonds and F13 protein

Figure 6 shows Valacyclovir’s 3D and 2D representations. The interaction between Valacyclovir and F13 (Figure 6) encompasses various non-covalent interactions, including hydrogen bonds, pi-alkyl interactions, and van der Waals forces.

**Fig 6.**
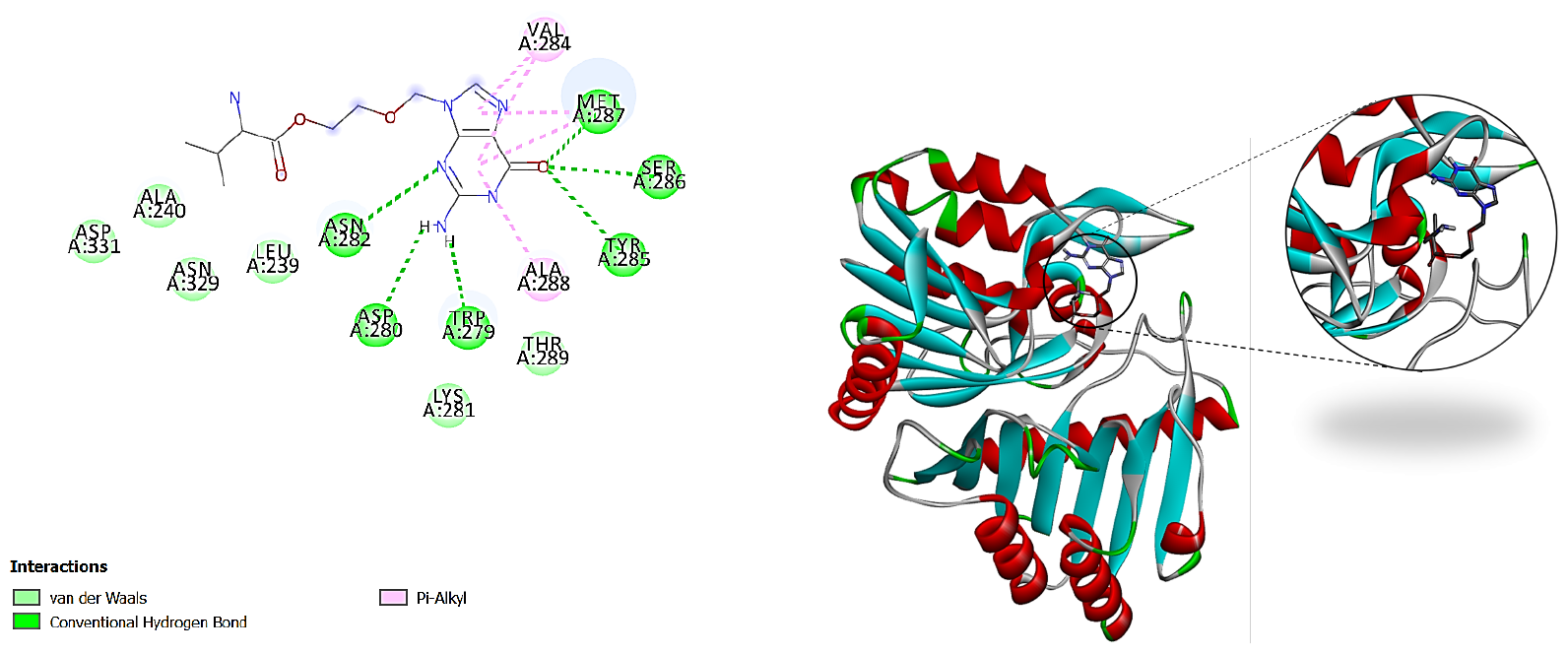
2D and 3D representations of the docked pose for the predicted interaction between Valacyclovir. The green dashed lines represent hydrogen bonds and F13 protein

Figure 7 shows Ribavirin’s 3D and 2D representations. This figure illustrates undesirable donor-donor bonding involving TYR285 and VAL284 in the interaction between ribavirin and F13. Nevertheless, the overall interaction pattern involves hydrogen bonds with TRP279, ASN282, MET287, and SER286 residues, along with van der Waals interactions with other residues.

**Fig 7.**
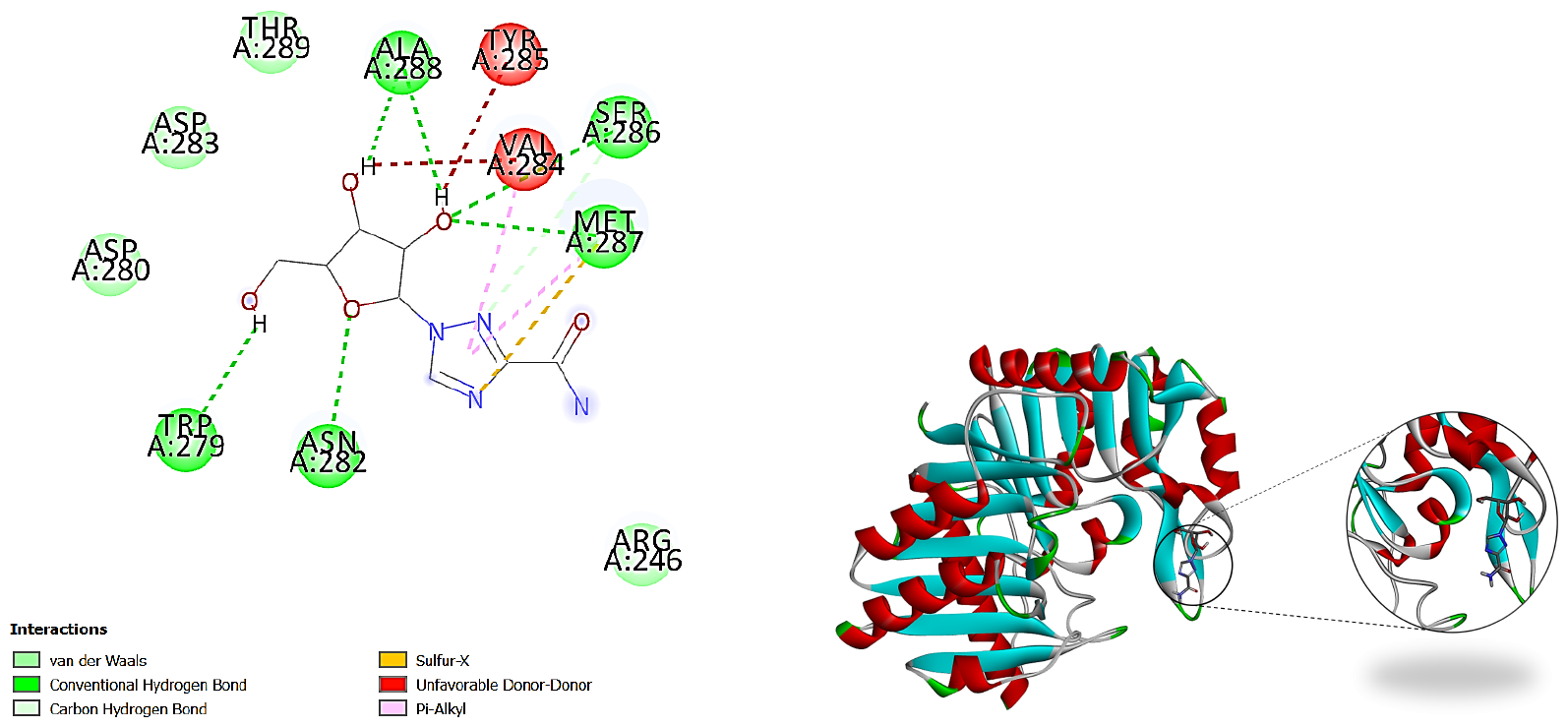
2D and 3D representations of the docked pose for the predicted interaction between Ribavirin. The green dashed lines represent hydrogen bonds and F13 protein

Figure 8 shows Favipiravir’s 3D and 2D representations. For the interaction of favipiravir with F13 (Figure 8), conventional hydrogen bonds, pi-alkyl interactions, and weak van der Waals bonds dominate.

**Fig 8.**
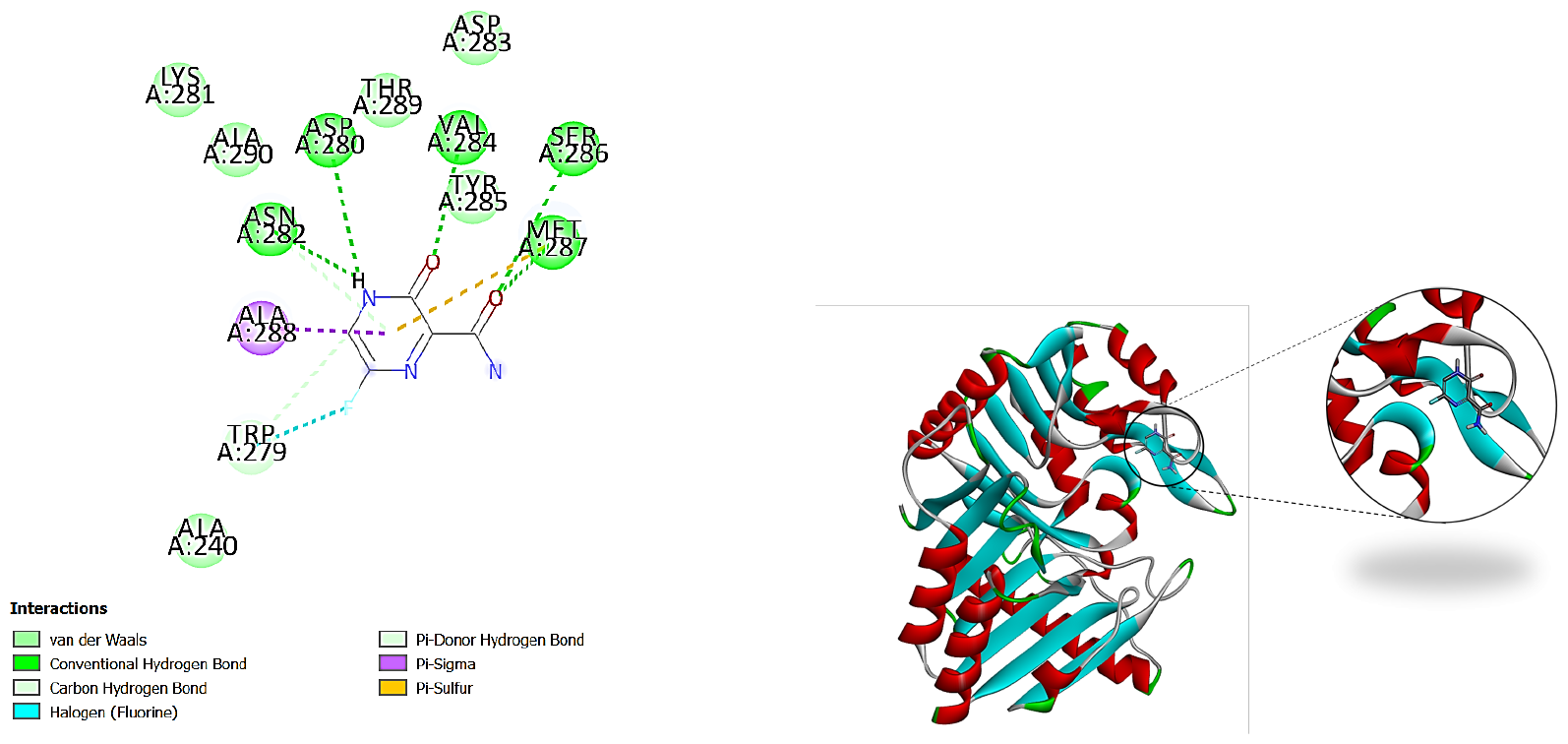
2D and 3D representations of the docked pose for the predicted interaction between Favipiravir. The green dashed lines represent hydrogen bonds and F13 protein

As the last suggested drug, Figure 9 shows Baloxavir marboxil’s 3D and 2D representations. Baloxavir binds to the F13 protein by forming hydrogen bonds with ASP280, VAL284, SER286, ASN282, and MET287 residues, as well as other interactions such as pi-sigma, pi-sulfur, and van der Waals forces.

**Fig 9.**
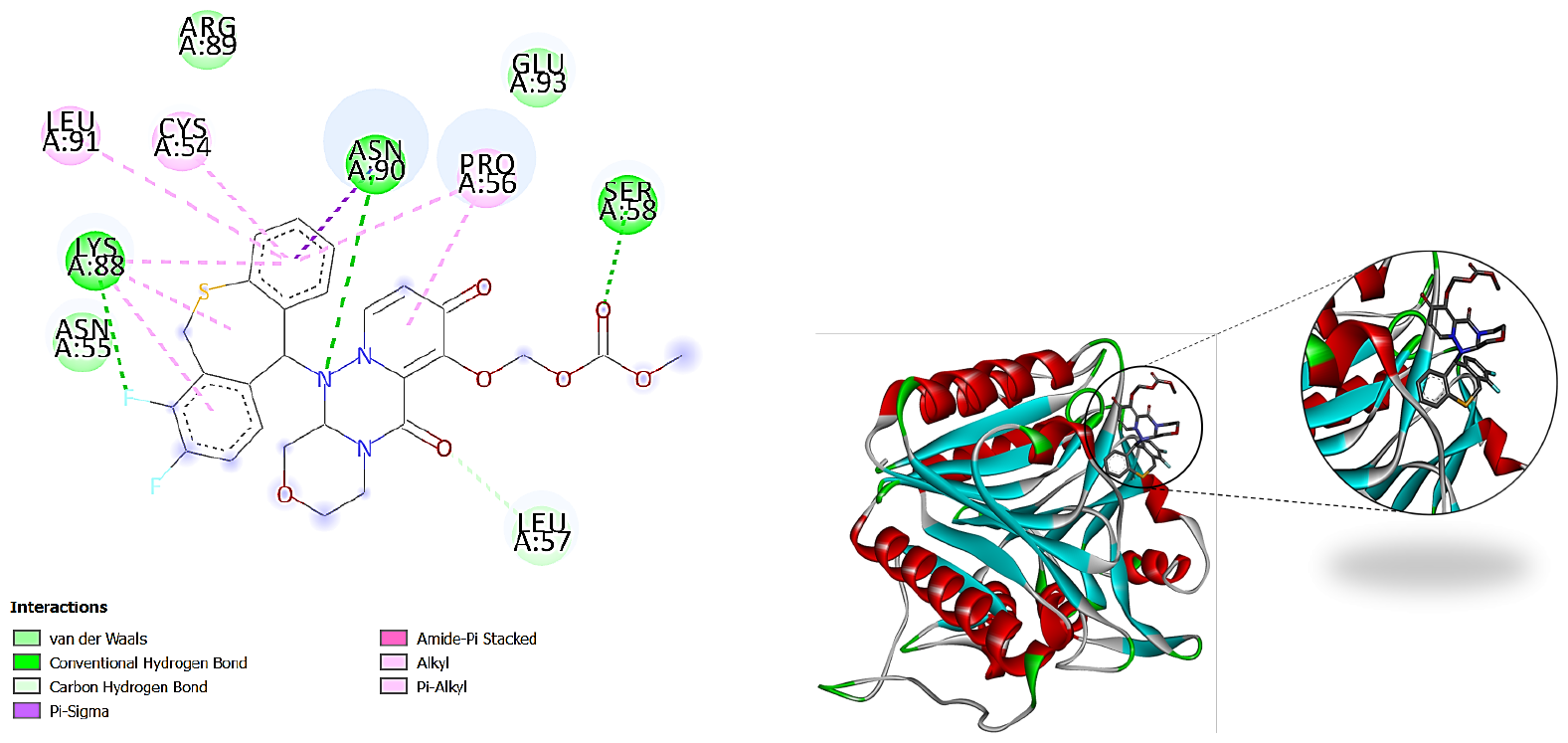
2D and 3D representations of the docked pose for the predicted interaction between Baloxavir marboxil. The green dashed lines represent hydrogen bonds and F13 protein

## 4 Conclusion

While the monkeypox virus has caused a global epidemic in recent years, there are no approved drugs for its treatment. This work proposed a computational drug repurposing approach to suggest several drugs to deal with MPXV. To do this, we prepared a dataset from viruses and antivirals to apply computational learning methods for drug suggestions. We have applied SVM, random forest, decision tree, and convolutional deep model (DRaW) to the virus and antiviral features to learn virus-antiviral interactions. The performance analysis shows that DRaW has the highest performance and the random forest comes after that. Then, using voting, the highly promising antivirals, i.e., Tilorone, Valacyclovir, Ribavirin, Baloxavir marboxil, and Favipiravir, suggested by the learning methods have been chosen for further analysis. We did the homology modeling and molecular docking study on the proposed drugs. The homology modeling using AlphaFold makes it clear that envelope receptor F13 can be the target of the antivirals. Therefore, we applied the docking studies on the suggested drugs of the computational modeling and it approved the hypothesis with high binding affinity. Generally, to our best knowledge, this is the first work that proposes antivirals for treating the MPXV. These screening results on Tilorone, Valacyclovir, Ribavirin, Baloxavir marboxil, and Faviapiravir can be further analyzed using laboratory analysis. The work approves the high potential of computational drug repurposing for the screening phase of drug discovery which causes lower time and costs.

## Availability of data and materials

The data and code of MPXV-Pred are freely available at https://github.com/BioinformaticsIASBS/MonkeyPox.

## Consent for publication

Not applicable.

## Authors’ contributions

MH and AZ conceptualized the idea. SMH implemented the methods. MHS performed the sequence alignment and prepared the drug similarity matrix. All authors participated in writing the manuscript. AZ prepared and wrote the docking studies. MH supervised and administered the project. MH and MHS read and approved the final manuscript.

